# A novel *ISG15* transcript restricts influenza A virus infection

**DOI:** 10.64898/2026.01.15.699784

**Authors:** Himadri Nath, Yan Guo, Steven F. Baker

## Abstract

The interferon response orchestrates antiviral defense through the induction of interferon-stimulated genes (ISGs). Although nearly all human genes undergo alternative splicing, the identity and function of many ISG transcript isoforms remains poorly understood. Using long-read RNA sequencing of human cells that were undergoing an interferon response, we identified a previously unrecognized interferon-inducible *ISG15* transcript. This non-canonical ISG15 (ncISG15) transcript contains a unique 5′ untranslated region and encodes an N-terminally truncated protein in frame with ISG15. During interferon stimulation, ncISG15 produces an ISGylation pattern distinct from canonical ISG15 and potently inhibits influenza A virus polymerase activity. The influenza A viral interferon antagonist protein NS1 counteracts ncISG15-mediated restriction; however, endogenous ncISG15 executes modest antiviral activity against NS1-encoding viruses. Antiviral activity of ncISG15 is largely independent of its function as a cytokine, through ISG augmentation, or by ISGylation. This novel transcript adds a new layer of regulatory complexity to the ISG15-driven antiviral network.

## INTRODUCTION

Cells rapidly respond to parasitic invasion to limit pathogen replication. Upon sensing viral components, cells produce type I interferons (IFNs) that trigger the induction of >1,000 interferon-stimulated genes (ISGs) that together create an antiviral state. ISGs have broad activities, including functioning as antiviral effectors that directly interfere with viral infection, signaling molecules that potentiate inflammation, and proteins that resolve inflammation to induce post-infection homeostasis^1^. A growing body of evidence suggests that beyond transcriptional induction, co- or post-transcriptional regulation, particularly alternative splicing, provides an additional layer of modularity in antiviral responses^2–4^. Conservative estimates in IFN-α treated B cells find that there are on average 2-3 transcripts per ISG^5^. These alternative isoforms often functionally negate their canonical isoform^6,7^. Previously, we have shown two host genes that through alternative splicing modulate influenza virus infection^8,9^. Despite this, the transcript isoform diversity of ISGs and their function in lung cells in the context of infection remains largely unexplored.

To identify alternatively spliced or otherwise hidden ISG transcripts, we used long-read RNA-seq (PacBio Iso-Seq). Short-read RNA-seq offers high depth and reliable quantification, but its fragmented reads make it difficult to reconstruct full-length isoforms, particularly those present at low abundance or with unusual splice junctions. Iso-Seq addresses this limitation by capturing complete transcript molecules and directly reports exon-intron sequences, alternative junctions, and atypical 5′ or 3′ configurations^10^. Although long-read sequencing produces data with comparatively low depth of coverage, its ability to resolve full-length isoforms makes it advantageous to discover novel transcripts. Using this approach, we detected an unannotated ISG15 transcript that retains most of the canonical open reading frame but differs at its 5′ end, consistent with an alternative splicing or alternative transcription initiation event. We refer to this transcript as non-canonical ISG15 (ncISG15).

ISG15 is one of the earliest and most strongly induced interferon-stimulated genes, reaching among the highest expression levels following IFN signaling or viral infection^11,12^. The protein comprises two ubiquitin-like domains connected by a short linker and is terminated with a conserved LRLRGG motif required for conjugation to lysine residues on target proteins. Conjugation of ISG15, termed ISGylation, is a post-translational modification catalyzed by the E1 activating enzyme UBA7 (UBE1L), the E2 conjugating enzyme UBE2L6 (UBCH8), and an E3 ligase (primarily HERC5). This dynamic process is reversed by the deISGylase USP18, generating a regulated post-translational modification cycle tightly coupled to IFN signaling as all five system components (ISG15, E1-E3, and USP18) are themselves ISGs^13–15^. ISG15 has three primary functions^16^. First, via protein ISGylation, it covalently modifies host and viral substrates to alter their stability, localization, or activity. ISGylation restricts diverse RNA and DNA viruses, including influenza A and B, Sindbis, vaccinia, and coxsackievirus by modifying viral proteins or essential host cofactors^16–18^. The influenza A NS1 protein, for example, is ISGylated by HERC5 at specific lysines, disrupting its dimerization and RNA-binding capacity^19^. Second, free intracellular ISG15 limits inflammation by stabilizing USP18. USP18 not only functions as the major deISGylase but also acts as a negative regulator of type I IFN signaling, preventing hyperactivation^20–22^. Third, secreted ISG15 acts as a cytokine-like molecule that promotes IFN-γ production from lymphocytes, linking innate antiviral defenses with adaptive immunity^23,24^. Together, these three modes of action underscore the pleiotropy of ISG15, functioning as both an effector and a regulator within the IFN network. ISG15 is also species- and virus-specific typified by 66% amino acid identity between mouse and man^25^. In mice, the absence of ISG15 or its conjugation machinery increases susceptibility to several viruses, whereas in humans, inherited ISG15 deficiency results in mycobacterial and not viral disease and an enhancement of basal IFN responses^23,26,27^. As a result of this specificity, human-adapted influenza B virus NS1 protein only binds to and inhibits primate ISG15s, alleviating viral restriction in human hosts^28–30^. The more promiscuous influenza A virus, however, remains sensitive to human ISGylation^31^.

Despite extensive knowledge on ISG15 function, ISG15’s transcript-level complexity remains ill defined. Our discovery of ncISG15 introduces a previously unappreciated dimension to ISG15 transcriptional and functional regulation. We observe that ncISG15 is expressed following IFN stimulation and viral infection and encodes an N-terminal truncated protein that exerts potent inhibition of influenza A virus polymerase activity. Mechanistically, this isoform retains the conserved conjugation motif but inhibits influenza largely through and ISGylation- and inflammation-independent manner. Influenza A virus NS1 reversed ISG15 and ncISG15-mediated polymerase inhibition in isolation, but we propose that stoichiometric abundance differences between the two isoforms leaves endogenous ncISG15 less susceptible to NS1 inhibition during infection. Thus, ncISG15 represents a naturally occurring, interferon-inducible ISG15 isoform that expands the functional diversity of the ISG15 system and reveals a new layer of host adaptation between influenza viruses and the interferon response.

## RESULTS

### Identification of the non-canonical ISG15 transcript

Long-read RNA-sequencing was performed on RNA extracted from human A549 cells mock-treated, stimulated with IFN-β, or infected with influenza A virus (WSN) and revealed that ∼5,446 full-length transcripts (∼42%) were shared between all three conditions, whereas 11-13% were unique to each condition **(Figure S1A)**. Of note, the limited overlap in transcripts in interferon vs infected conditions is likely the result of limited depth of sequencing – better resolution could be obtained through more sequencing or by applying matched short-read RNA-seq data^32^. The majority of transcripts were present in RefSeq (full splice match), but ∼1/3 were novel (incomplete splice match, novel not in catalog) **(Figure S1B)**. Within the ISG15 locus, three distinct transcript isoforms were detected **(Figure S1C)**. Transcript 1 corresponded to the canonical two-exon ISG15 mRNA (RefSeq ID NM_005101.4; Ensembl ISG15-203). Transcript 2 represented a previously unannotated splice variant that lacked exon 1 and instead contained a non-canonical (nc) exon embedded within intron 1, then spliced into the shared exon 2 of canonical ISG15. We hereafter term this transcript ncISG15. A third isoform is unlikely to encode a stable protein due to a very small open reading frame, and it was not pursued further. Of note, we did not detect the other three annotated transcripts from Ensembl (ISG15-201, 202, 204), which could be explained by limited depth in our sequencing. Junction-spanning short-read RNA-sequencing analysis from our prior data confirmed ncISG15 splicing and suggested levels similar to other antiviral ISGs *APOBEC3G, ETV7*, and *ZBP1* **(Figure S1D)**^9^.

To further characterize the abundance and regulation of the ncISG15 transcript, we designed isoform-specific primers and TaqMan probes **(Figure 1A)**. Each transcript was amplified using a common reverse primer in exon 2 and unique forward primers recognizing exon 1 or the nc exon sequences. End-point RT-PCR from oligo-dT-primed cDNAs confirmed robust amplification of both canonical and ncISG15 mRNAs in IFN-stimulated and virus-infected A549 cells, while PB2 amplicons verified infection and MECR served as a housekeeping control **(Figure 1B)**. Comparable amplification patterns were observed in human bronchial epithelial (HBEC) cells immortalized from four different donors^33,34^ **(Figure 1C)**. TaqMan quantitation showed that ISG15 is expressed at basal levels of ∼10^2^ copies, and induced 100-fold to ∼10^5^ copies **(Figure 1D)**. By contrast, ncISG15 was undetectable without stimulation, but induced 1,000-fold to ∼10^3^ copies. ISG15 and ncISG15 copy numbers in HBECs varied between individuals but followed the trend of A549 cells (**Figure S1E**). These findings establish ncISG15 as a bona fide interferon-stimulated transcript.

**Figure 1.**
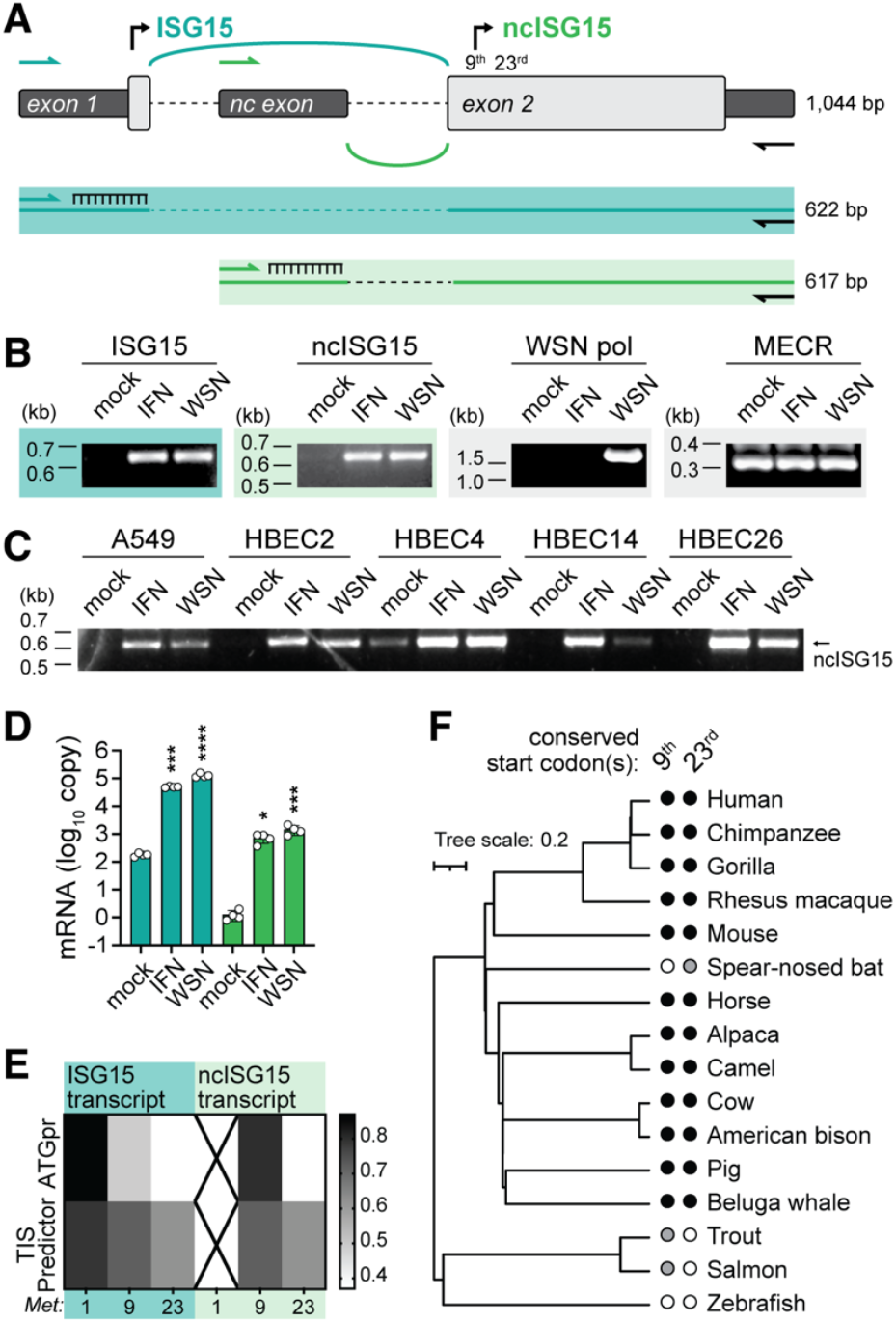
Identification, expression, and conserved predicted translation of ncISG15. (A) Schematic of the *ISG15* locus highlighting canonical exon 1, the nc exon, and shared exon 2. Locations of isoform-specific amplification strategies (primers and probes) and expected base pair (bp) product sizes are indicated. (B) RT-PCR of ISG15 and ncISG15 transcripts in A549 cells during mock or IFN-α treatment (100 U/ml; 18 h), or influenza A virus infection (WSN; MOI, 0.1; 18 h). Viral polymerase (WSN pol) and MECR amplified as controls. (C) Detection of ncISG15 across A549 and human bronchial epithelial cell (HBEC) lines from different donors, treated as in (B). (D) Absolute mRNA abundance of ISG15 and ncISG15 (log_10_ copy number) from A549 cells, treated as in (B). (E) Predicted translation initiation strength across candidate methionines in ISG15 and ncISG15 transcripts using ATGpr and TIS Predictor. (F) Phylogenetic tree of ISG15 orthologs including presence (black filled circle) or absence (empty circle) of start codons at the 9^th^ and 23^rd^ positions. Gray filled circle indicates start codon 1-2 positions away. Data in panel D represent mean ± SD of technical replicates from two independent biological experiments. Statistical comparisons were performed using one-way ANOVA with Dunnett’s multiple comparisons test relative to mock; *, *P* < 0.05; ***, *P* < 0.001; ****, *P* < 0.0001.

To assess its translation potential, we analyzed Kozak sequences in ncISG15 using ATGpr and TIS Predictor algorithms^35–37^. Both tools yielded similar results – minimal leaky initiation at the 9^th^ codon from the canonical transcript, but robust initiation at the same codon in the nc transcript **(Figure 1E)**. This codon is in frame with the canonical ISG15 ORF, producing an 8 amino acid (MGWDLTVK) N-terminally truncated version of ISG15. Comparative alignment across mammals revealed that this methionine is highly conserved, with analogous residues observed in vertebrates **(Figure 1F)**. Considering ISG15’s relatively low amino acid identity across species, conservation of the 9^th^ methionine may suggest evolutionary preservation of the alternative translation product. Together, these results validate ncISG15 as an interferon-inducible transcript capable of producing an N-terminally truncated product.

### ncISG15 inhibits influenza A polymerase activity more potently than ISG15

Prior studies demonstrated that ISG15 limits influenza virus multiplication^18,29,31^, though its role on influenza A polymerase (3pol) activity has yet to be assessed. We thus performed minigenome assays in 293T cells expressing canonical and ncISG15 and observed a dose-dependent decrease of 3pol activity with increasing expression of either ISG15 isoform, with a more pronounced reduction for ncISG15 **(Figure 2A)**. Western blotting showed ncISG15 accumulates to lower abundance compared to ISG15 despite equimolar plasmid concentrations. However, at the higher concentration of ncISG15, there is reduced abundance of the viral nucleoprotein (NP), supporting a model in which the alternative isoform impairs viral polymerase function by decreasing NP levels. Together, these observations reveal that ncISG15 exerts potent antiviral activity against influenza A virus replication despite its limited accumulation.

**Figure 2.**
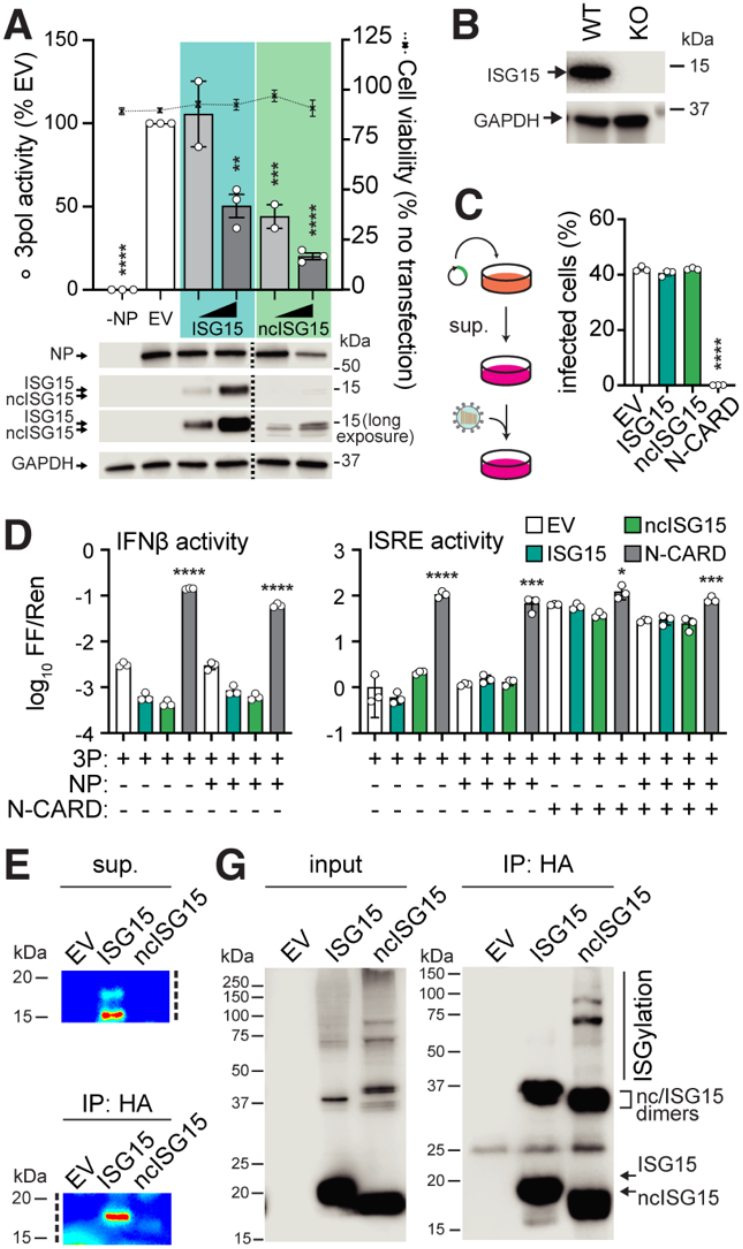
ncISG15 inhibits influenza A polymerase activity and displays distinct ISGylation patterns. (A) Influenza A virus polymerase (3pol) activity in 293T cells in the absence of NP (-NP) or with NP plus empty vector (EV), ISG15, or ncISG15. Cell viability is indicated on the right axis. Representative immunoblots are shown below. (B) Immunoblot of ISG15 accumulation in wild-type (WT) and ISG15 knockout (KO) cells treated with 250 U/ml IFN-□ for 18 h. (C) Conditioned media from 293T cells expressing EV, ISG15, ncISG15, or RIG-I N-CARD was transferred (supernatant, sup.) onto A549GFP1-10 cells prior to WSN PA-2A-11x7 infection (MOI, 0.1; 6 h). (D) IFN-β and ISRE promoter activity in 293T ISG15-KO cells expressing ISG15, ncISG15, or N-CARD. (E) Detection of secreted HA-tagged proteins from 293T ISG15-KO cells expressing HA-ISG15 or HA-ncISG15 directly from supernatant or after immunoprecipitation (IP). (F) Anti-HA IP of ISG15- or ncISG15-associated proteins, showing ISGylated species and isoform-specific dimers. (G) Data in A represent grand mean from 2-3 independent experiments ± SEM normalized to EV. C, and D are representative mean ± SD data from at least three independent experiments. Statistical significance was determined using one-way ANOVA with Tukey’s multiple comparisons test relative to EV controls; *, P < 0.05; **, P < 0.01; ***, P < 0.001; ****, P < 0.0001.

### Functional characterization of ncISG15: secretion, signaling, and ISGylation biology

Given their similarity, we sought to evaluate whether ncISG15 functions in a similar manner as the three main roles of ISG15: (1) secretion into the extracellular milieu, (2) free intracellular signaling that can modulate interferon responses, and (3) covalent conjugation to target proteins through ISGylation. To evaluate ncISG15 function in isolation we generated ISG15-knockout (KO) cells using CRISPR-Cas9 **(Figure 2B)**. We first used a bioassay to test whether ncISG15 secretion or its expression induces a secreted molecule that can inhibit influenza virus infection in recipient cells (**Figure 2C**). Whereas the positive control N-CARD, a constitutive activator of MAVS and subsequent cytokine production, completely abolished infectivity in recipient cells, neither ISG15 nor ncISG15 expression inhibited infection. We next tested whether free intracellular ncISG15 influences interferon signaling by transfecting KO cells in the absence of IFN (and thus E1-E3 expression) **(Figure 2D)**. In the presence of inactive or active 3pol, IFN-β promoter and interferon-stimulated response element (ISRE) reporter activity showed no significant difference among cells expressing canonical or ncISG15. By contrast, N-CARD expression strongly induced both reporters, but under these stimulated conditions we did not observe ISG15-mediated reduction in ISRE activity. Prior data showed that free ISG15 reduces IFN extracellular signaling, which we observe slightly for both ISG15 and ncISG15 in the IFN reporter assay in the absence of N-CARD^27^. Both canonical and ncISG15 isoforms were detected in immunopurified culture supernatants, demonstrating that ncISG15, like ISG15, is secreted, albeit to lower levels corresponding with intracellular abundance **(Figure 2E)**. These findings indicate that ncISG15, although secreted, likely does not inhibit influenza polymerase activity through an IFN-mediated manner.

We finally turned to the third major mode of ISG15 action, protein ISGylation. Covalent conjugation of ISG15 to lysine residues on target proteins requires induction of the IFN-stimulated E1-E3 enzymes UBE1L, UBCH8, and HERC5. To evaluate if ncISG15 could similarly be conjugated onto target proteins we analyzed lysates of transfected and IFN treated, E1-E3 expressing, or influenza-infected A549 cells by western blotting against ISG15 to query for the ladder-like ISGylation pattern. While both isoforms produced an ISGylation pattern, ncISG15 displayed a distinct pattern, with a subset of higher molecular weight species (**Figure 2F**). The presence of unique bands in the ncISG15 lanes suggests that the ncISG15 engages a partially divergent set of host substrates, or that there is differential partial degradation of proteins modified with ISG15 *vs* ncISG15. Proteins modified by canonical or ncISG15 could be immunopurified, which further confirmed the presence of distinctly modified proteins as well as the characteristic ISG15 or ncISG15 dimer. Of note, we also pulled down canonical or ncISG15 in the context of a minigenome assay followed by blotting against PB2, PB1, PA, and NP, but did not observe modified viral proteins (data not shown).

### Reduction in viral polymerase activity by ncISG15 is largely ISGylation-independent

Prior work has shown that ISGylation is a major mechanism limiting both influenza A and B virus growth^18,29^. To evaluate whether the antiviral activity of ncISG15 is primarily mediated through ISGylation, we performed experiments designed to interrogate this pathway (**Figure 3A**). First, we ablated canonical or ncISG15 conjugation capability by mutating the terminal amino acids from GG (wildtype) to AA and performed polymerase activity assays in ISG15-KO 293T cells. For the remainder of polymerase activity assays, we used equimolar, low plasmid amounts of ISG15 and ncISG15 that moderately or severely, respectively, restrict polymerase function. Polymerase activity both in the presence or absence of IFN did not differ significantly between the GG and AA variants for either isoform **(Figure 3B-C)**. We next co-expressed E1-E3 ISGylation enzymes to increase ISGylation and observed that canonical ISG15 + E1-E3 more significantly decrease polymerase activity compared to ISG15 alone, whereas the reduction mediated by ncISG15 + E1-E3 was modest but not statistically 6 significant **(Figure 3D)**. Lastly, we induced ISGylation via IFN treatment and co-expressed the deISGylase UPS18 to remove conjugates and find similar results as above – USP18 rescued ISG15-mediated polymerase inhibition but had a modest and non-significant effect on ncISG15 **(Figure 3E)**. Collectively, these results suggest that the antiviral function of ncISG15, in contrast to ISG15, occurs largely through an ISGylation-independent manner.

**Figure 3.**
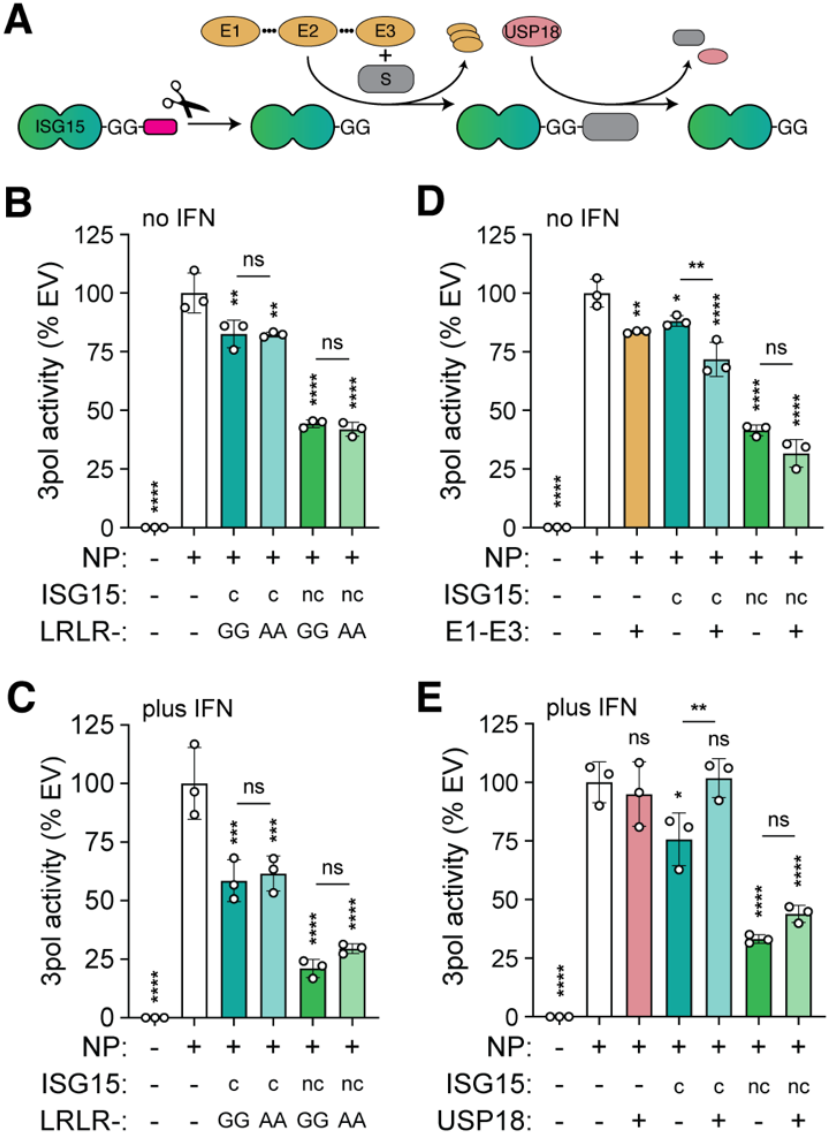
ncISG15-mediated restriction of influenza polymerase activity is largely ISGylation-independent. (A) Schematic of the ISGylation pathway and experimental perturbations. (B-E) Influenza polymerase (3pol) activity in cells expressing canonical (c) or non-canonical (nc) ISG15. (B-C) Polymerase activity with wildtype C-terminal diglycine (GG) or conjugation-deficient (AA) motifs in the absence (B) or presence (C) of 250 U/ml IFN. (D) Polymerase activity in the presence of co-expressed ISGylation enzymes (E1-E3: UBA7, UBE2L6, and HERC5) without IFN treatment. (E) Polymerase activity following IFN-□ treatment (250 U/ml, 6 h post transfection) with or without the addition of the deISGylase USP18. (F) Data are shown normalized to EV control and are representative mean ± SD from two-three independent experiments with technical replicates. Statistical comparisons were performed using one-way ANOVA with Tukey and Dunnett’s multiple comparisons test; ns, not significant; *, P < 0.05; **, P < 0.01; ***, P < 0.001; ****, P < 0.0001.

### ncISG15 restricts influenza A virus

As artificial overabundance of ncISG15 limited polymerase activity, we next assessed whether transient expression of ncISG15 could similarly limit whole virus replication. ISG15-KO 293T cells expressing ISG15, ncISG15, or as a positive control the anti-influenza factor IFITM3, were infected with a NanoLuc luciferase-expressing influenza A virus, and virus growth was quantified by titrating supernatants on fresh cells^38^. Expression of ncISG15, but not ISG15, resulted in a significant reduction in viral titers **(Figure 4A)**. The ∼50% reduction in titer by ncISG15 was similar to, but less significant than that of IFITM3 overexpression. As in polymerase activity assays, equimolar amounts of ISG15 and ncISG15 were used for transfection, thus it is possible that modest ISG15-mediated polymerase restriction is not observable in infection settings. We also tested the effect of overexpression on adenovirus C5 E1-like nano-luciferase gene expression, but in contrast to the antiviral brincidofovir, these antiviral genes had no effect^39,40^. These findings indicate that ncISG15 overexpression restricts influenza A virus replication.

**Figure 4.**
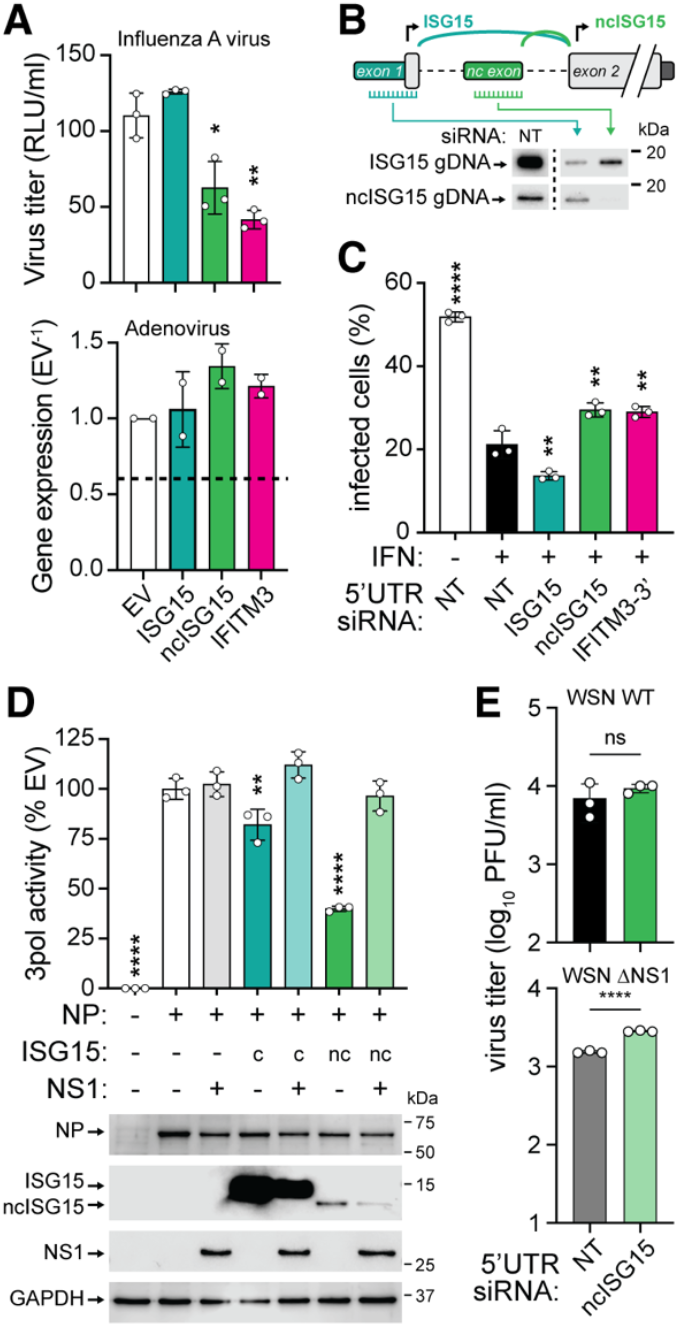
ncISG15 restricts influenza A virus replication and is antagonized by NS1. (A) Top, influenza A virus (WSN NLuc) production from 293T ISG15-KO cells expressing EV, ISG15, ncISG15, or IFITM3 (MOI, 0.01; 24 h). Bottom, human adenovirus (HAdV-C5 NLuc) reporter expression in cells (MOI, 0.1, 24 h), dotted line indicates NLuc expression with 25 µM brincidofovir. (B) Schematic of isoform-specific siRNA targeting strategies for ISG15 and ncISG15. (C) Validation of isoform-specific knockdown using genomic DNA constructs (gDNA) by immunoblot. (D) Flow cytometry-based evaluation of influenza A virus (WSN PA-2A-11x7) infection during transcript-specific knockdown (MOI, 0.1, 6 h). (E) Influenza polymerase (3pol) activity in 293T ISG15-KO cells in the presence of canonical (c) or non-canonical (nc) ISG15 with or without influenza A virus NS1. Representative immunoblots shown below. (F) Viral titers of wildtype or ΔNS1 influenza A virus (WSN; MOI, 0.1; 24 h) following infection of cells with ncISG15 knockdown. (G) Data are representative mean ± SD from three independent experiments with technical replicates. Statistical analysis was performed using one-way ANOVA with Dunnett’s multiple-comparisons test. ns, not significant; *, P < 0.05; **, P < 0.01; ***, P < 0.001; ****, P < 0.0001.

We next sought to evaluate the contribution of endogenous ncISG15 to antiviral defense, which is made difficult due to the bicistronic nature of *ISG15*. We designed custom siRNAs that targeted *ISG15* exon 1 or the nc exon to selectively degrade ISG15 or ncISG15 transcripts, respectively, and report here one siRNA targeting each isoform **(Figure 4B)**. The siRNAs were validated in ISG15-KO cells expressing ISG15 or ncISG15 genomic constructs, with western blot analysis showing isoform-specific depletion. We next assessed how endogenous *ISG15* products impact influenza A virus infection. A549 cells transfected with each siRNA were treated with IFN-α to induce *ISG15* expression, then subsequently infected with our GFP-expressing influenza A virus **(Figure 4D)**^41^. Flow-cytometric quantification of GFP-positive cells revealed that IFN treatment in non-targeting siRNA transfected cells reduced viral infection from 50% to 20%. ISG15 selective knockdown resulted in a more pronounced IFN-mediated antiviral phenotype, likely through enhanced IFN signaling due to the loss of ISG15’s USP18-stabilizing function^27^. In contrast, depletion of ncISG15 partially restored virus infectivity, comparable to the effect observed upon knockdown of IFITM3, suggesting that endogenous ncISG15 activated by IFN contributes to anti-influenza host defense.

### Influenza A NS1 protein counteracts ncISG15-mediated antiviral activity

Due to the fact that ncISG15 exerts antiviral activity, it is likely that influenza A viruses would be evolutionarily selected for a trait that counteracts this function. The influenza A non-structural protein 1 (NS1) is a multifunctional interferon antagonist that suppresses host antiviral defenses by targeting innate immune signaling, host mRNA processing, and interferon-stimulated effector pathways^42,43^. For example, NS1 inhibits RIG-I mediated signaling and interferon induction through protein:protein interactions with host ubiquitin ligases and RNA sensors^44^. We thus evaluated whether influenza A virus NS1 counteracts the anti-influenza activity of ncISG15. Strikingly, co-expression of NS1 restored polymerase activity suppressed by either ISG15 isoform, indicating that NS1 effectively reverses their inhibitory effects **(Figure 4E)**. This further suggests that both canonical and ncISG15 restrict polymerase activity at a step susceptible to NS1-mediated antagonism. Western blot analysis showed that both ISG15 and ncISG15 accumulation is moderately reduced in the presence of NS1 suggesting NS1 transcriptionally or post-translationally regulates canonical and ncISG15 accumulation. Despite NS1 antagonism, transient expression of ncISG15 significantly reduced virus titers **(Figure 4A)**. We then infected IFN-treated ncISG15-knockdown cells with wildtype or ΔNS1 virus and measured virus growth by plaque assay **(Figure 4F)**. While there was a modest but non-significant growth advantage for wildtype virus in the absence *versus* presence of ncISG15, ΔNS1 virus had a significant growth advantage when endogenous ncISG15 was knocked down. This indicates that viruses lacking NS1 are hypersensitive to ncISG15-mediated restriction. These findings demonstrate that influenza A virus NS1 protein specifically counteracts the antiviral activity of endogenously produced ncISG15, revealing a previously unrecognized interface between viral immune evasion and ISG15 transcript-variant regulation.

## DISCUSSION

Here we identify a previously unrecognized interferon-induced *ISG15* transcript encoding an isoform that subverts influenza A virus replication. This non-canonical ISG15 isoform, revealed through long-read RNA-sequencing, lacks canonical exon 1 but contains a non-canonical exon within ISG15’s only intron. ncISG15 likely originates from alternative transcription initiation, although no additional ISRE promoters were found between exon 1 and the nc exon. Because we did not experimentally distinguish between alternative initiation or splicing, we report only that ncISG15 is a novel ISG15 transcript. The ncISG15 transcript was observed by long- and short-read RNA-seq, and by RT-PCR amplification from human lung epithelial cell lines and from cells derived from four different human donors. Custom RT-qPCR showed that ncISG15 is induced by three log_10_ copies from IFN treatment or influenza viral infection compared to mock treatment, while ISG15 is induced by two log_10_ copies. However, the total pool of transcripts during stimulation contains ∼two log_10_ copies more ISG15 than ncISG15. The ncISG15 transcript encodes a protein translated from a highly conserved internal methionine, producing an in-frame truncated version of ISG15 lacking the first eight amino acids. Plasmid-based expression of ncISG15 with an N-terminal HA tag accumulates less than ISG15. This difference is not due to transcript stability or proteasomal or lysosomal turnover (data not shown). Like ISG15, ncISG15 also likely ISGylates proteins, but showed a unique ISGylation signature. Additionally, ncISG15 was detected in cell supernatants. These observations indicate preserved induction, modification, and secretion, suggesting that ncISG15 is a naturally regulated component of the ISG15 system rather than an aberrant transcript.

Functionally, ncISG15 exhibited stronger antiviral activity against influenza A virus than canonical ISG15. Both isoforms reduced polymerase activity, but the inhibition was markedly greater for ncISG15. Yet, when viral replication was measured in overexpression or isoform-specific knockdown experiments, only ncISG15, and not ISG15, significantly restricted virus infection. We propose a logical explanation for this observation: ncISG15 escapes NS1 antagonism due to expression differences. During infection-induced expression of both canonical and ncISG15, the majority of NS1 will be sponged by ISG15 due to the two log_10_ expression difference in ISG15 vs ncISG15. This interpretation is further supported by the observation that silencing ncISG15 enhanced ΔNS1 virus replication. Western blot data showed that ncISG15 overexpression reduced NP accumulation, and that NS1 overexpression reduced ISG15 and ncISG15 accumulation. Future experiments will determine the mechanism of restriction and its antagonism by NS1.

These findings also add context to several earlier findings on ISG15 that appeared inconsistent across systems. Studies in human cells showed that ISGylation is antiviral against influenza A virus and it can modify influenza A NS1 and reduce NS1 activity^18,31^. Work in mice demonstrated that loss of ISG15 leads to pronounced viral susceptibility, consistent with a strong antiviral function^17^. In contrast, analyses of humans with inherited ISG15 deficiency revealed normal or even enhanced resistance to many viral infections due to an elevated interferon response, which arises because ISG15 is required to stabilize USP18 and consequently repress IFN receptor signaling in humans^45^. Notably, this regulatory relationship differs in mice, where USP18 stability does not depend on ISG15, explaining the stronger antiviral phenotype observed in knockout mice^45^. Our results suggest that ncISG15 reconciles these observations: canonical ISG15 is readily neutralized by influenza NS1 in human cells, whereas ncISG15 retains antiviral activity and is less susceptible to NS1-mediated antagonism. Thus, the anti-influenza effects attributed to *ISG15* in humans may predominantly reflect ncISG15, while canonical ISG15 functions chiefly as a regulator of interferon homeostasis.

By analyzing the known functions of ISG15, we find that ncISG15’s antiviral mechanism operates largely independent of ISGylation or by augmenting the interferon response. Whereas ISG15 antiviral activity was enhanced by overexpressing the E1-E3 ISGylation machinery and reversed in the presence of the deISGylase USP18, ncISG15 antiviral function in these conditions was modest but not significantly affected. Given that most antiviral functions of canonical ISG15 depend on conjugation to host or viral proteins, our evidence suggests ncISG15 may act through non-covalent interactions. Our experiments with promoter reporters or with conditioned media show that ncISG15 does not restrict virus by inducing or augmenting the antiviral state. Failed attempts to co-immunoprecipitate ncISG15 with viral polymerase components motivates us to in the future explore non-covalent host protein interactions.

Several observations from an evolutionary perspective reinforce the role of ncISG15 in limiting viral pathogenesis. First, despite ISG15’s low sequence identity across species, the 9^th^ methionine is highly conserved. It remains to be tested whether other vertebrates deploy ncISG15 and whether this occurs through a transcriptionally dependent mechanism, as with humans, or through another manner. Second, the fact that influenza A virus NS1 antagonizes ncISG15 function highlights an evolutionary balance between host antiviral defense and viral evasion. This presents an interesting challenge for the virus – non-specific destruction of ISG15, in an attempt to destroy ncISG15, could have the unintended consequence of enhancing IFN signaling due to decreased USP18. The functional difference between ISG15 and ncISG15 could be explained by different protein:protein interactions, which NS1 may be less able to differentiate. It will be important to identify the precise targets of ncISG15 and to determine whether NS1 physically engages with ncISG15. Third, we found that ncISG15 does not restrict influenza B virus polymerase activity (data not shown). Moreover, influenza B NS1 does not antagonize ncISG15-mediated restriction of influenza A virus polymerase activity despite its ability to antagonize ISG15-mediated restriction of influenza B NP (data not shown)^29^. The observation that influenza B NS1 cannot counteract ncISG15-mediated antiviral activity of influenza A virus fits a co-evolutionary model where influenza B virus insensitivity to ncISG15 provided a background where evolution of an ncISG15 escape mechanism was irrelevant. The specificity could reflect structural differences in the viral polymerase complex or divergent influenza A and B NS1s. Such selectivity suggests that the interferon system may exploit alternative isoforms like ncISG15 to fine-tune antiviral breadth across virus families.

In conclusion, we describe ncISG15 as a novel, interferon-inducible *ISG15* transcript that exerts potent antiviral activity against influenza A virus that is functionally antagonized by NS1. The finding that a single ISG can produce isoforms with distinct regulatory and antiviral properties underscores the complexity of innate immune regulation. Prior functional descriptions of ISG alternative transcripts have shown a preponderance for IFN-suppression that resolves inflammation coordinated by the canonical isoform^4^. Given the prevalence of alternative splicing among ISGs, additional uncharacterized isoforms exist that may, like ncISG15, have specialized roles in host defense. Understanding how these variants interact with viral antagonists and whether unique regulatory systems control their expression will provide additional insight into the molecular arms race between host and pathogen and may guide the design of antiviral strategies that exploit natural isoform diversity.

## ACKNOWLEDGEMENTS

We thank members of the Baker lab for vibrant discussion and critical reading of the manuscript. This work was supported by the American Lung Association (ERPALA2023).

## AUTHOR CONTRIBUTIONS

Conceptualization: HN, SFB; methodology: HN, SFB; formal analysis, HN, YG, SFB; investigation, HN, YG, SFB; writing – original draft, HN & SFB; writing – review & editing, HN, YG, SFB; visualization, SFB; funding acquisition, SFB; supervision, SFB

## METHODS

### Cell lines

A549, 293T, MDCK, MDCK-TMPRSS2^46^ cells were maintained in Dulbecco’s Modified Eagle Medium (DMEM; Gibco) supplemented with 10% fetal bovine serum (FBS) and incubated at 37°C with 5% CO2. A549^GFP1-10^ cells fluoresce upon infection with a GFP11-encoding virus^41^. Human bronchial epithelial cells (HBECs) from 4 different donors were maintained in Keratinocyte serum-free media (Gibco) on FNC-coated plastic (Athena)^33,34^. Cells were routinely evaluated for mycoplasma (Lonza). For viral infections, virus growth medium (VGM) was used, consisting of DMEM supplemented with 0.2% bovine serum albumin (BSA), 25 mM HEPES, 1x penicillin-streptomycin, and TPCK-treated trypsin (0.25–2 μg/mL; Sigma).

ISG15-deficient A549 and HEK293T cells were generated using a lentiviral CRISPR-Cas9 system targeting the human ISG15 locus. Guide RNA targeting exon 2 was synthesized in the lentiCRISPRv2 backbone (GenScript). Lentivirus was produced in HEK293T cells by co-transfecting the lentiCRISPRv2-ISG15 plasmid with packaging vectors pMD2.G and psPAX2 (Addgene #s 12259 and 12260), and clarified viral supernatants were used to transduce target cells in the presence of 8 µg/ml polybrene. Transduced cells were selected with puromycin for 5-7 days until untransduced control cells were fully eliminated. Knockout efficiency in clonal cells was assessed by Sanger sequencing of the targeted genomic locus followed by analysis using Synthego’s ICE (Inference of CRISPR Edits) tool and subsequently confirmed by immunoblotting for ISG15 under basal and IFN-stimulated conditions.

### Viruses

Influenza A viruses were generated by reverse genetics using strain A/WSN/1933 (H1N1; WSN). 293T cells were transfected with eight pTM or pBD viral genome plasmids (PB2, PB1, PA, HA, NP, NA, M, NS) and supporting expression plasmids (p3X-1T, pCAGGS NP, TMPRSS2) using jetPRIME (Polyplus) one day prior to adding MDCK-TMPRSS2 cells as previously described^47–49^. Reporter influenza viruses contained bicistronic PA segments additionally encoding NanoLuc (NLuc) or GFP11x7^38,41^. Rescued viruses were plaque purified on MDCK cells, sequence-confirmed by Oxford Nanopore (Plasmidsaurus), and titrated by plaque assay on MDCK cells. WSN virus lacking NS1 (ΔNS1; kindly provided my A. Mehle) was titered on MDCK cells overexpressing NS1-GFP^50^. Human species C5 adenovirus encoding NLuc upstream of E1A was provided by C. King and previously described^39^.

### Plasmid constructs

Plasmids used in this study were generated using standard molecular cloning approaches, including Gibson assembly (New England Biolabs), inverse PCR-based mutagenesis, and Golden Gate assembly, following established principles from our prior work^9^. Coding sequences for canonical ISG15 and the non-canonical ISG15 isoform were cloned into pCMV-HA-IFITM3 kindly provided by J. S. Yount^51^. CRISPR-immune ISG15/ncISG15 constructs were prepared using inverse-PCR to introduce silent mutations in the ISG15 gRNA target sequence. Conjugation-deficient nc/ISG15 mutants were generated by substituting the C-terminal LRLRGG motif with LRLRAA through site-directed mutagenesis. Genomic constructs of ISG15 and ncISG15 were cloned from genomic DNA: pcDNA3 g-ISG15 contains the entire *ISG15* gene with 5’UTR and 3’UTR, pcDNA3 g-ncISG15 is the same but lacks the 5’UTR and intron 1 until the nc exon. E1 activating enzyme pCAGGS-HA-hUBE1L (UBA7), and E2 conjugating enzyme pFlagCMV2-UbcH8 (UBE2L6) were provided by Addgene (#s 12438 & 12442). E3 ligase pcDNA6.2 HERC5-V5 was assembled from pENTR223 (DNASU HsCD00516476) using Clonase II (Thermo), and pcDNA3 V5-USP18 ORF was synthesized (Twist) and cloned into pcDNA3 by Gibson assembly. Constitutively stimulatory form of RIG-I (pEF-Tak-FLAG RIG-I (1-228); termed N-CARD) was kindly provided by A. M. Kell^52^. Influenza pcDNA3 V5-NS1A (WSN) and V5-NS1B (B/Brisbane/60/2008) were cloned from pBD and pHW2000 plasmids, respectively^53^. pCAGGS B/Yamagata/16/1988 PB2, PB1, PA, NP, and pPolI B-HA-luc were described previously^54^. All constructs were sequence-verified prior to use (Plasmidsaurus).

### Long-read RNA sequencing

Iso-Seq (PacBio) was performed to obtain full-length transcript resolution. Total RNA was extracted from A549 cells left untreated (mock), stimulated with IFN-α (250 U/mL for 8 h), or infected with influenza WSN (MOI, 0.02; 24 h)^9^. TRIzol-extracted RNA was quality verified (RIN > 8.5), and three biologic RNA replicates per condition were pooled prior to library construction (University of Wisconsin-Madison Biotechnology Center). Full-length cDNA was synthesized from poly(A)+ RNA using the Iso-Seq Express Oligo-dT protocol (PacBio). SMRTbell libraries were generated, size-selected to enrich for full-length products, and sequenced on the PacBio Sequel II platform across two SMRT cells. Raw subreads were processed and aligned to the human genome (hg38) using the PacBio Iso-Seq3 pipeline^55,56^. To complement long-read sequencing, matched samples (mock, IFN-β treated, influenza-infected) were processed for short-read RNA-seq (BioProject PRJNA667475). Libraries were prepared for Illumina sequencing and aligned to hg38^9^.

### Manual inspection and comparative visualization of long- and short-read data

Iso-Seq full-length reads and short-reads (bam and indexed bam files) from mock-, IFN-β–treated, and virus-infected samples were visualized using Integrated Genome Viewer (IGV). Candidate unannotated isoforms were identified through manual inspection of (i) full-length read clusters with noncanonical exon-intron organization, (ii) splice junctions lacking existing annotations, and (iii) reproducible 5′ or 3′ transcript extensions across conditions and sequencing modalities. Novel transcript structures were shortlisted when supported by both long- and short-read RNA-seq data. Exon 1-2 junction spanning reads of ISG15, ncISG15, APOBEC3G, ETV7, ZBP1, and MECR was performed in IGV (sashimi function) across 3 biological replicates of short-read data.

### RT-PCR and RT-qPCR validation of ISG15 transcripts

Transcript-specific forward primers targeting exon 1 or the nc exon, along with a common exon 2 reverse primer, were designed for end point RT-PCR. RNA was extracted from A549 or HBEC cells and cDNA prepared using oligoDT. One Taq hot start DNA polymerase (NEB) was used for PCR.

For RT-qPCR, specific forward primers and probes was used for each transcript. The reverse primer was the same for both transcripts. Prime time one step RT-qPCR mix was used, and products analyzed on Quantstudio 5. Custom primers and probes (IDT).

### In silico ORF analysis

Putative translation initiation sites within candidate ISG transcripts were evaluated using two independent computational tools, ATG-PR and TIS-Predictor, which score potential start codons based on local nucleotide context, Kozak consensus strength, predicted ribosome accessibility, and comparative feature models^36,37^. For ISG15 or ncISG15 transcripts, the full-length cDNA sequence was provided as input, and all in-frame methionines were interrogated. ATG-PR generated ranked initiation-site probabilities derived from logistic regression models trained on experimentally validated initiation events, whereas TIS-Predictor applied a deep-learning–based framework incorporating codon neighborhood and sequence motifs. The first, ninth, and 23^rd^ codon were analyzed (ISG15 numbering) and resulting likelihood of transcription initiation was predicted where higher scores indicate higher likelihood.

### Phylogenetic analysis

ISG15 ortholog amino acid sequences of representative species were retrieved from NCBI as follows: Human, NP_005092.1; Chimpanzee, XP_054956204.1; Gorilla, XP_055237821.1; Rhesus macaque, NP_001253735.1; Mouse, NP_056598.2; Spear-nosed bat, XP_028368350.2; Horse, XP_005607662.1; Alpaca, XP_015093270.1; Camel, XP_014406705.1; Cow, NP_776791.1 ; American bison, XP_010848324.1; Pig, NP_001121941.2; Beluga whale, XP_030619807.1; Trout, XP_029618957.1; Salmon, NP_001134702.1; Zebra fish, NP_001191098.1. Sequences were aligned using CLUSTAL MUSCLE (3.8) with default parameters. Alignments were imported into Figtree for visualization, manual refinement, and annotation. Trees additionally annotated with the presence or absence of ATG start codons at the 9^th^ or 23^rd^ position, or the 16^th^ (salmon and trout) or 24^th^ (spear-nosed bat) positions.

### Polymerase activity assays

293T WT or ISG15-KO cells were seeded in 24-well plates and transfected in triplicate with pCAGGS or p3X1T plasmids encoding PB1, PB2, and PA, and pCAGGS NP or empty vector using TransIT-X2 (Mirus). ISG15, ncISG15, NS1, E1-E3, and USP18 plasmids were co-transfected as specified. A firefly luciferase reporter plasmid (pHH21-vNA-Luc), flanked by influenza virus UTRs, was co-transfected to assess viral polymerase activity (PMID: 18692771). A plasmid constitutively expressing Renilla luciferase served as a normalization control (pRL-SV40, Promega). Twenty-four hours post-transfection, cells were lysed, and firefly and Renilla luciferase activity was measured. Lysates were also analyzed by western blot to verify expression transfected constructs using the following antibodies: rabbit anti-PB2 (PMID: 19995968); rabbit anti-PA, GeneTex GTX125933; goat anti-NP, BEI NR-3133; rat anti-GFP, Chromotek 3H9; and rabbit anti-β-actin, GeneTex GTX637675.

### Cell viability assay

Cell viability was measured using the CellTiter-Glo® 2.0 Cell Viability Assay (Promega). Briefly, cells were seeded in opaque-walled 96-well plates and transfected with polymerase activity assay conditions as indicated. Control wells containing culture medium without cells were included to subtract background luminescence. After 48 h, plates were equilibrated to room temperature and assayed per manufacturer’s protocol. Luminescence values were normalized to a no transfection control.

### Conditioned media bioassay

293T ISG15-KO cells were transfected with empty vector, HA-ISG15, HA-ncISG15, or N-CARD. After 24 h, culture supernatants were collected, clarified by centrifugation (500 x *g* for 5 min), and diluted 1:1 in fresh medium. The diluted supernatants were added to A549^GFP1-10^ cells seeded the day prior. Following 24 h of conditioning, cells were infected with WSN PA-2A-11x7 virus. At 6 h post-infection, cells were detached using Trypsin-EDTA, fixed with 3% paraformaldehyde, and analyzed by flow cytometry to quantify GFP-positive cells.

### IFN-β and ISRE activity assays

293T ISG15-KO cells were seeded in 24-well plates and transfected in triplicate with 3X1T (PB2, PB1, PA), pRL SV40, pcDNA3 mCherry, and pHH21 vNA-GFP, and additionally with empty vector or pCAGGS NP to prevent or activate influenza polymerase activity observed by GFP production. Cells were additionally transfected with ISG15 or control constructs and ISRE FF-Luc (pHISG54-Luc) or IFN**-**β FF-Luc (pIFΔ(−116)lucter) plasmids to quantify innate immune stimulation^57,58^ (kindly provided by L. Martínez-Sobrido and A. J. te Velthuis). 24 h later, firefly luciferase activity normalized to Renilla luciferase was measured from cell lysates.

### Immunoprecipitation

For detection of ISGylation patterns, ISG15 KO 293T cells were transfected with HA-ISG15 or HA-ncISG15 constructs. Eight h post transfection; cells were treated with 1,000 U/ml IFN-α for 18 additional hours. Cells were then lysed in ice-cold Co-IP lysis buffer (50 mM Tris pH (7.4), 150 mM NaCl, 0.5% NP-40 substitute, protease inhibitor cocktail), clarified by centrifugation, and used immediately for immunoprecipitation (IP). For secretion experiments, culture supernatants were collected 48 h post-transfection, clarified, and then used for IP. First, the lysate or supernatant was precleared with agarose beads (Santa Cruz SC-2001), then nutated overnight at 4°C with anti-HA (10 µg/ml; clone 3F10, MilliporeSigma). Next day the lysate antibody mix was affinity purified using Dynabeads (Thermo, 10003D). To preserve ISGylation, eluates or inputs were resolved by SDS-PAGE under non-reducing, non-boiled conditions and visualized by western blotting with anti-ISG15 (Proteintech, 15981-1-AP).

### Viral infection during transient expression of ISG15 or ncISG15

ISG15-KO 293T cells were reverse-transfected with plasmids expressing HA-ISG15, HA-ncISG15, HA-IFITM3, or empty vector using Transit X2 in 96-well plates. Twenty-four h post-transfection, cells were infected with WSN NLuc (MOI, 0.01). 24 h post-infection, culture supernatants were collected and used for virus titration. Viral titers were determined by infecting MDCK cells for a single replication cycle (9 h), after which NLuc activity was measured using Nano-Glo luciferase assay system (Promega).

Adenovirus (HAdV-C5) infections were performed in parallel using the same transient-transfection as above in ISG15-KO 293T cells. Cells transfected with empty vector plus or minus brincidofovir (25 µM)^59^, HA-ISG15, HA-ncISG15, or IFITM3 were infected with HAdV-C5-NLuc (MOI, 0.1; 24 h) and luciferase activity was measured directly from infected cells.

### Validation of selective of ISG15 transcript siRNAs

siRNAs were custom-designed to specifically target exon 1 of ISG15 or the nc exon of ncISG15 (Horizon ON-TARGETplus). To validate the knockdown efficiency, 293T ISG15 KO cells were reverse transfected with 50 nM siRNA in a 24-well plate for 24 h. Cell media was then replaced with fresh media and cells were transfected with 62.5 ng ISG15 or 125 ng ncISG15 genomic constructs. Whole-cell lysates were collected 24 h later and assessed by western blotting with an anti-ISG15 antibody.

### Influenza A virus infection during transcript-specific knockdown

#### Flow cytometry-based single-cycle infection assays

A549^GFP1-10^ cells seeded in a 24-well plate were transfected with 50 nM siRNAs targeting ISG15, ncISG15, IFITM3 (Horizon, L-014116-01), or a non-targeting (NT) control (Horizon, D-001810-10) using Transit X2. After 24 h of siRNA treatment, cells were stimulated with 100 U/mL IFN-α for 24 h. Cells were then infected with WSN PA-2A-11x7 (MOI, 0.1). At 6 h post-infection, cells were detached with Trypsin-EDTA, fixed with 3% PFA and analyzed by flow cytometry to quantify GFP-positive infected cells.

#### Plaque assays to determine multicycle impact

A549 cells were transfected with siRNAs against ISG15, ncISG15, IFITM3, or NT control as described above. After 24 h, cells were treated with 100 U/mL IFN-α for 24 h and subsequently infected with either WSN WT or ΔNS1 (MOI, 0.1). At 24 h post-infection, culture supernatants were collected and titrated on MDCK cells using a standard plaque assay.

## SUPPLEMENTAL INFORMATION

**Supplemental Figure 1.**
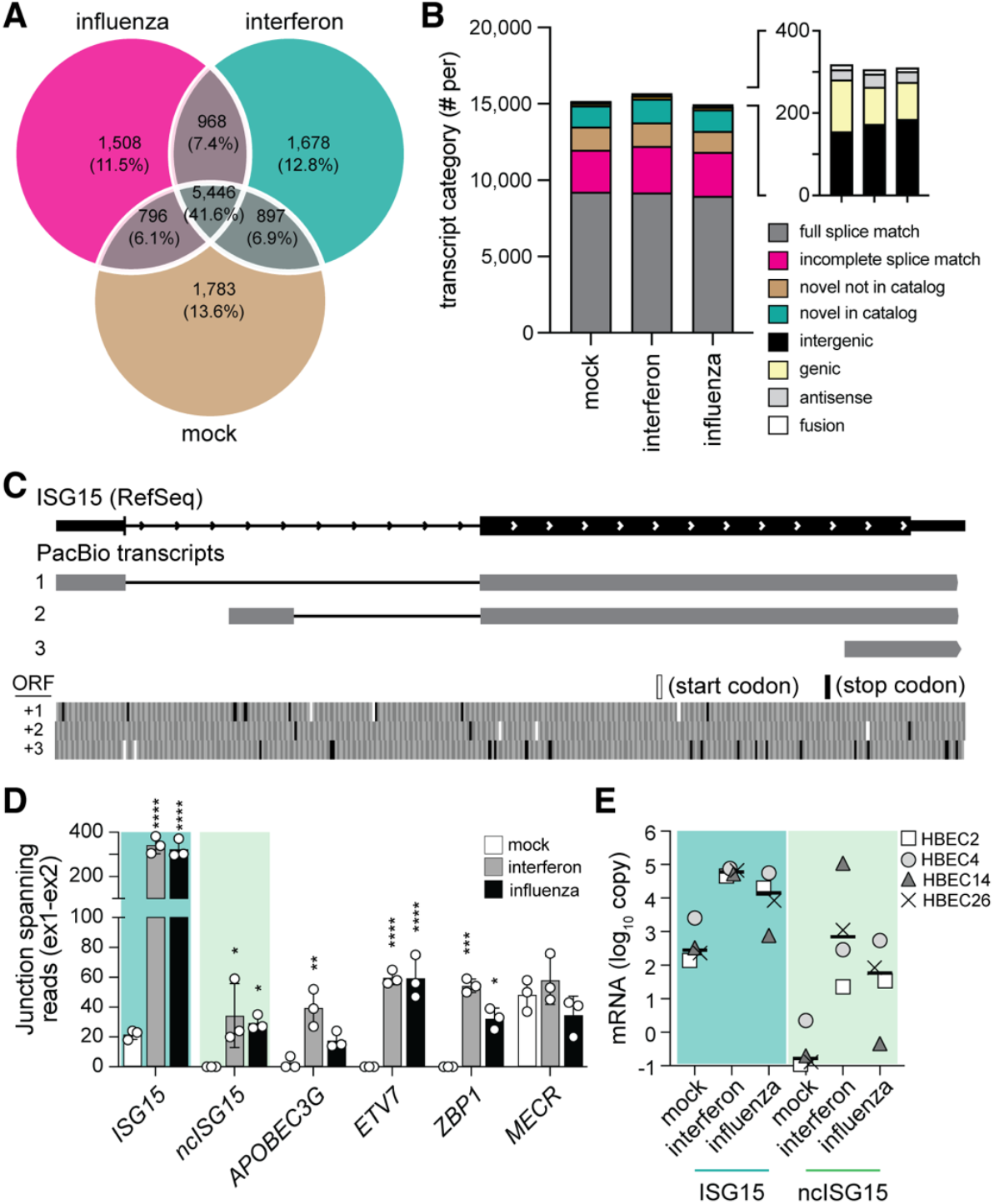
Long-read RNA-sequencing reveals ncISG15. (A-B) Overview of long-read RNA-seq transcriptomics. (A) Overlap of full-length transcripts detected across mock or IFN treatment, or influenza A virus infection. (B) Distribution of transcript categories identified by Iso-Seq. (C) ISG15 locus showing transcripts detected by long-read RNA-seq (PacBio) with indicated open reading frames (ORFs 1-3) and potential start/stop codons. (D) Junction-spanning read counts (exon 1-2) from short-read RNA-seq (3 biologic replicates; BioProject PRJNA667475) for ISG15, ncISG15 and selected antiviral or control genes in mock, IFN, and influenza conditions. (E) ISG15 and ncISG15 absolute mRNA abundance (log10 copy number) determined by RT-qPCR from HBEC lines during mock, IFN-□, and influenza conditions (treated as in Fig. 1). Statistical comparisons in panel D were performed using one-way ANOVA with Dunnett’s test relative to mock. *, P < 0.05; **, P < 0.01; ***, P < 0.001; ****, P < 0.0001.

